# Genetic selection for small molecule production in competitive microfluidic droplets

**DOI:** 10.1101/469007

**Authors:** Larry J. Millet, Jessica M. Velez, Joshua K. Michener

## Abstract

Biosensors can be used to screen or select for small molecule production in engineered microbes. However, mutations to the biosensor that interfere with accurate signal transduction are common, producing an excess of false positives. Strategies have been developed to avoid this limitation by physically separating the production pathway and biosensor, but these approaches have only been applied to screens, not selections. We have developed a novel biosensor-mediated selection strategy using competition between co-cultured bacteria. When applied to biosynthesis of *cis,cis*-muconate, we show that this strategy yields a selective advantage to producer strains that outweighs the costs of production. By encapsulating the competitive co-cultures into microfluidic droplets, we successfully enriched for muconate-producing strains from a large population of control non-producers. Facile selections for small molecule production will increase testing throughput for engineered microbes and allow for the rapid optimization of novel metabolic pathways.

## Introduction

Engineered microbes can produce a range of valuable bioproducts, providing the basis for a renewable bioeconomy^1^. However, initial strain construction efforts are often inefficient, and the cost and effort required to optimize a proof-of-concept pathway can preclude further development. New strategies for constructing libraries of mutant strains have accelerated the design and build phases of the engineering cycle^2–4^, but these efforts are limited by the throughput of methods for identifying improved variants. Demonstrations of new library construction methods frequently use phenotypes that can be linked to microbial growth or to production of an easily-identifiable molecule, such as visibly-pigmented compounds. Using similar approaches to engineer strains that produce higher concentrations of an arbitrary small molecule is challenging.

One option for identifying rare variants from a mutant library is to include a biosensor, such as a metabolite-responsive transcription factor or regulatory RNA, that reports on production of the target small molecule^5,6^. These biosensors can be highly effective, but they also come with several limitations. Most notably, the biosensor is typically expressed from the same cell that is being engineered or evolved. As a result, mutations can occur in the sensor that disrupt the desired regulation, producing false positive signals that require significant additional screening^6^. An intracellular biosensor also reports on the intracellular small molecule concentration. A product that is rapidly exported may not show a differential signal as the production rate changes. Similarly, a product that is rapidly imported may produce crosstalk between neighboring cells. Finally, biosensors can be linked to selectable markers such as antibiotic resistance genes or screenable markers such as fluorescent proteins. Selections typically enable larger library sizes and require less infrastructure, but also amplify any concerns about false positives during the selection.

Several strategies have been proposed to overcome these challenges using biosensors to screen mutant libraries. Avoiding false positives requires separating the biosensor from the strain being engineered in order to limit mutation accumulation in the biosensor. After the two functions are separated, though, measurements of productivity then require either disruptive methods to reintroduce the sensor into the producer strains^7^ or some form of spatial structure to prevent crosstalk between members of the library. Initial efforts often used colonies spread on petri dishes, which limited the assay throughput^8^. More recent approaches rely on encapsulation of cells inside microfluidic droplets^9–12^. By physically separating individual cells into distinct droplets, interactions between cells can be eliminated. However, all of these approaches depend on quantitative screens, limiting throughput and requiring fluorescence-activated cell sorters or on-chip droplet sorters to isolate desired droplets. A biosensor-driven selection strategy would simplify the process of isolating improved variants from a mutant library while expanding the potential library size.

We have developed a selection strategy that uses competition between bacterial strains within microfluidic droplets to select for small molecule biosynthesis. Production of the target small molecule inhibits growth of a sensitive competitor and provides a selective advantage to the producer (Figure 1A). We show that competitor strains can be rationally engineered to be inhibited by a target small molecule, that production of the target provides an advantage to the producer, and that spatial structure is required to prevent interactions between producer cells. By combining these elements, we enriched true producer strains from a background of faster-growing non-producers. These high-through selections for small molecule production will enable rapid optimization of engineered microbes.

**Figure 1:**
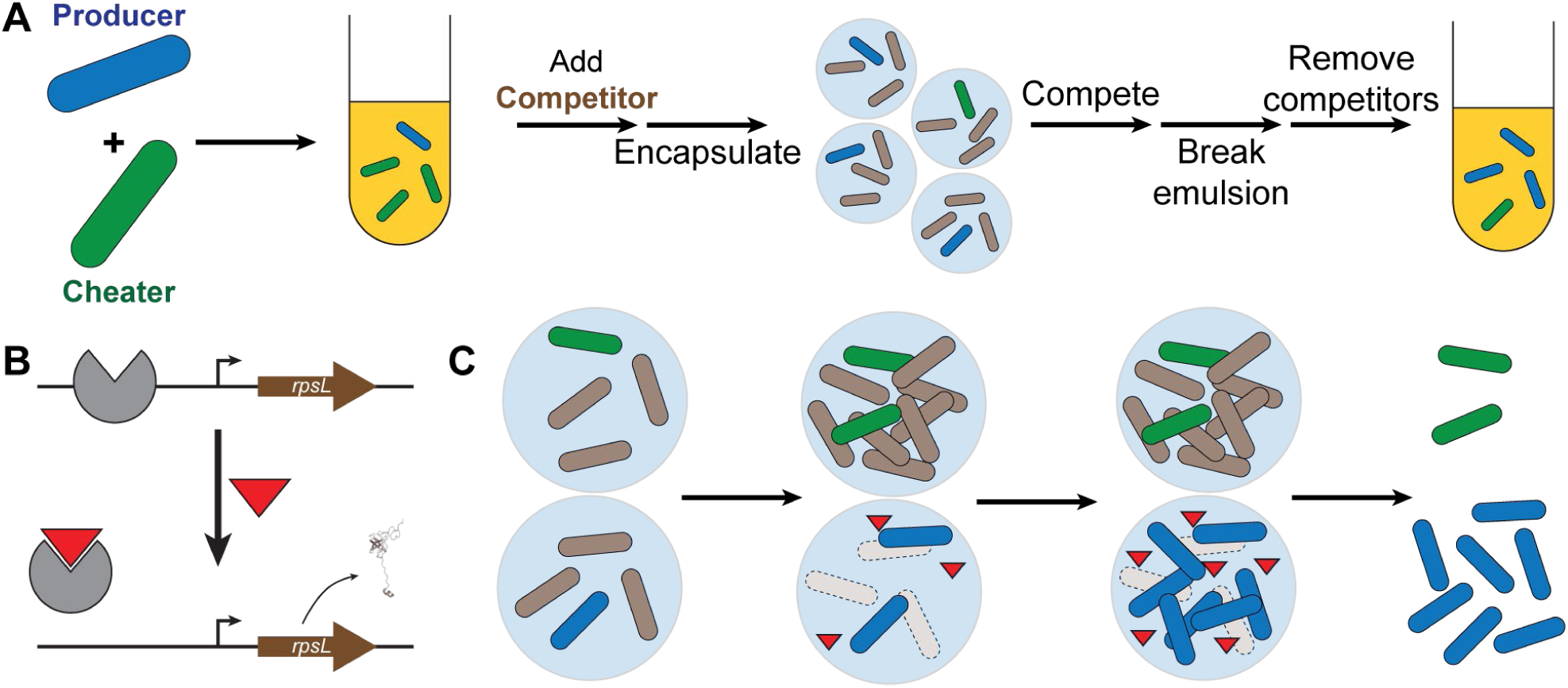
Competitive microfluidic droplets enable selections for small molecule production. (A) simplified population contains a mix of producer strains (blue) that synthesize the desired small molecule and cheater strains (green) that do not. This population is then combined with the competitor strain (brown) and encapsulated into microfluidic droplets where the competitor is inhibited by the target molecule. The mixed cultures are grown to saturation, the emulsion is broken, and the producer cells increase in frequency due to the fitness benefit of eliminating the competitor strain. (B) The competitor strain is inhibited by the target molecule (red triangle) due to regulation of the counterselectable marker *rpsL* by the biosensor. In the presence of the target molecule, production of RpsL sensitizes the competitor strain to streptomycin. (C) In a droplet containing a cheater, the competitor strain outcompetes the cheater and limits the yield. Conversely, in a droplet containing a producer, production of the target molecule blocks competitor growth and increases the yield of the producer strain.

## Results and Discussion

### Strain design and construction

To establish a microbial competition mediated by small molecule production, two different strains are inoculated into microfluidic droplets. The producer strain synthesizes a target small molecule, while the competitor strain is inhibited by the synthesized target (Figure 1). We chose muconate as our target small molecule, with *Pseudomonas putida* as the producer strain and *Escherichia coli* as the competitor.

The selection system consists of two strains, one engineered to produce muconate and the other engineered to be inhibited by muconate. To build a suitable muconate-producing strain, we started with *P. putida* CJ102, which was been previously engineered to synthesize muconate from lignin-derived aromatic compounds^13^. The strain was further modified by deletion of the adhesin *lapA*, to prevent biofilm formation that otherwise interfered with independent segregation^14^; deletion of a component of the type 6 secretion system *tssA1*, to eliminate an alternative killing mechanism^15^; deletion of the glucose dehydrogenase *gcd*, to prevent accumulation of a potential cross-feeding metabolite^16,17^; and introduction of a streptomycin-resistance marker^18^. The resulting strain, JMP15, was then further modified by deletion of the phenylpropanoid-CoA synthetase *fcs*^19^ to produce a control strain, JMP26, that is incapable of converting phenylpropanoids into muconate (Supplementary Figure 1A).

For the muconate-inhibited strain, we introduced a counterselection module into a streptomycin-resistant strain of *E. coli*. The counterselection module consists of a constitutive promoter driving expression of an operon containing a muconate transporter and a muconate-responsive transcription factor, combined with a regulated promoter driving expression of a counterselectable marker (Supplementary Figure 1B). We selected the *mucK* muconate transporter from *Pseudomonas putida* and the muconate-responsive transcription factor *benM* from *Acinetobacter baylyi*^20^, and then expressed a streptomycin-sensitive allele of *rpsL* from the BenM promoter. The counterselection module was synthesized *de novo* and cloned into pET-9a. When the resulting plasmid was transformed into streptomycin-resistant *E. coli* REL606, addition of muconate inhibited growth in a dose-dependent fashion, with growth completely blocked at 10 mg/L muconate (Figure 2).

**Figure 2:**
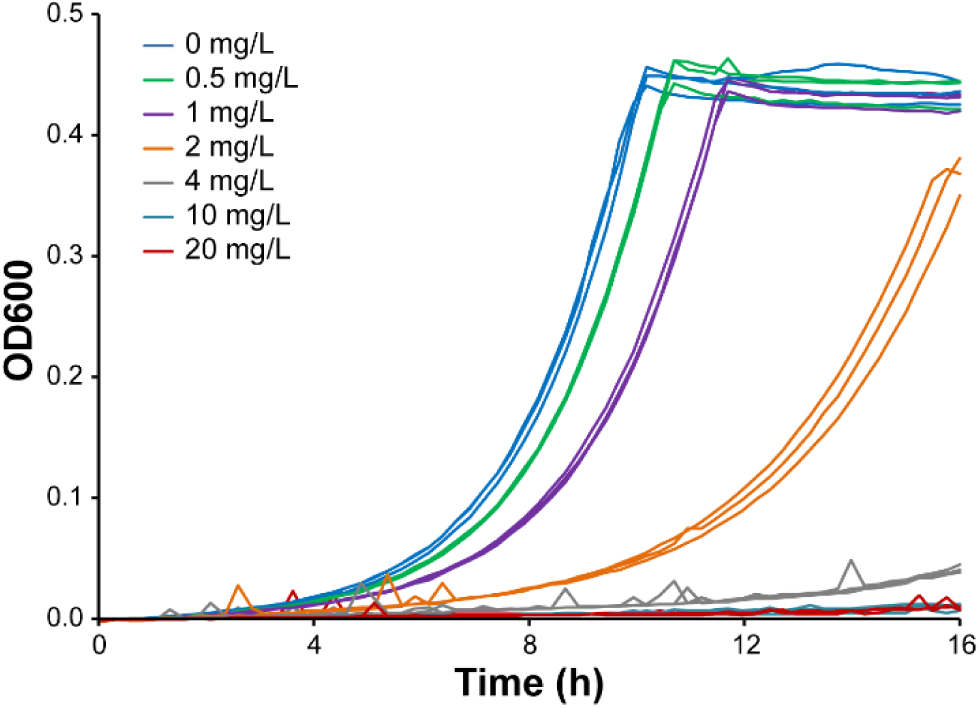
A muconate-responsive counterselection module demonstrates dose-dependent growth inhibition. Streptomycin-resistant REL606 containing plasmid pJM242 was grown in minimal medium containing ampicillin, streptomycin, and the indicated concentration of muconate. Three biological replicates are shown.

### Muconate production is costly but provides a competitive advantage against sensitive *E. coli*

The engineered producer strain, JMP15, can convert coumarate to muconate in a series of reactions that conserve little usable energy. Consequently, the strain grows more slowly in minimal media containing glucose when coumarate is also added (Figure 3A). The control strain, JMP26, does not show a growth difference.

**Figure 3:**
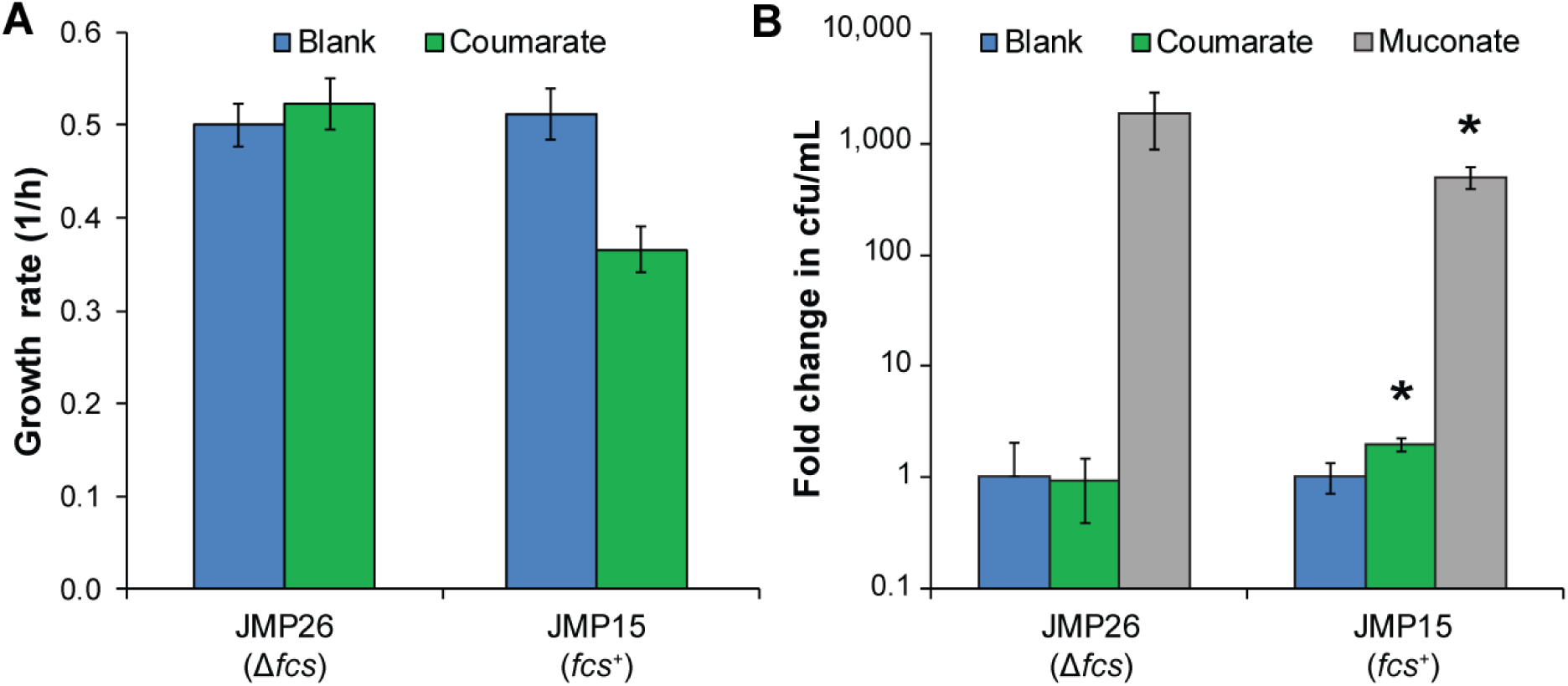
The benefits of muconate production outweigh the costs. (A) Addition of coumarate, which can be converted into muconate, decreases the growth rate of JMP15. Deletion of the gene encoding the first enzyme in coumarate metabolism, the CoA synthetase *fcs*, eliminates this effect. (B) When grown in competition with *E. coli* that are sensitive to muconate, the addition of coumarate provides a selective advantage only for the strain able to convert it into muconate. *: *p* < 0.05, two-tailed t-test. Error bars show one standard deviation, calculated from three biological replicates.

We then grew the producer and control strains of *P. putida* in co-culture with the muconate-sensitive *E. coli*. After growth, the *P. putida* concentration was measured by plating on selective agar. When exogenous muconate was added, both *P. putida* strains showed large increases in yield (Figure 3B). After addition of coumarate, the muconate-producing strain JMP15 showed a small but significant fold increase in yield of 1.9±0.2 while the yield of the control strain was unchanged. When grown in competition with a muconate-sensitive *E. coli*, muconate production is costly to the cell, but the benefits of inhibiting *E. coli* growth offset the costs.

### Selection of muconate producers from mixed cultures requires spatial structure

Muconate production provides an advantage when a pure culture of *P. putida* producers is competed against *E. coli* (Figure 3B). However, when using a mixture of producer and control *P. putida*, the costs are borne by the individual producers but the benefits, in terms of reduced *E. coli* competition, are shared. Therefore, when a mixture of control and producer *P. putida* are competed against *E. coli* in test tubes, we would expect the control strains to be more fit than the producers. We tested this hypothesis by growing all three strains together, and then separating out the *P. putida* and counting the fraction of JMP15 and JMP26 colonies. As predicted, the population fraction of JMP15 was unchanged, averaging a 1.0-fold enrichment (Figure 4A). Under these conditions, muconate production still increases the fitness of the *P. putida*, but the control strains share equally in the benefits.

**Figure 4:**
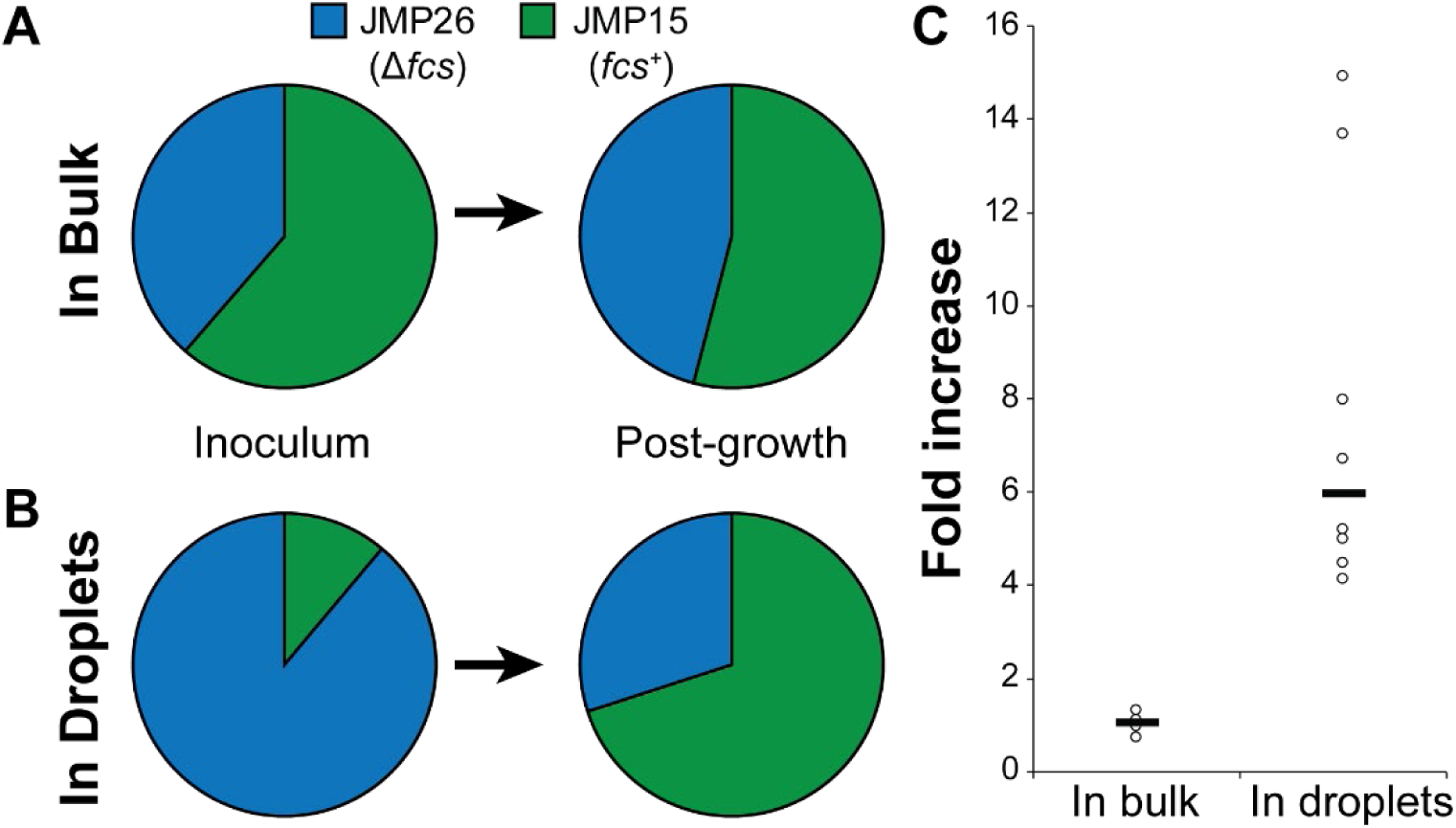
Spatial structure enables muconate selection. (A) A mixture of control and producer strains were competed against muconate-sensitive *E. coli* in test tubes. The fraction of JMP15 in the population did not change significantly. (B) When the mixed cultures were encapsulated into microfluidic droplets, a single night of growth enriched for muconate producers. One representative population ratio is shown for each condition. (C) The fold increase in JMP15 relative to JMP26 was calculated for each competition culture. Each point represents a single biological replicate, and each set of competitions was repeated on at least two days. White circles: individual cultures. Black bars: Distribution median.

Successful selection for efficient producers, therefore, requires a method to limit the benefits of muconate production to the producers. Microfluidic droplets can provide the necessary spatial structure, by encapsulating each producer cell into a separate droplet. When a similar three-way competition was conducted in droplets, the JMP15 population ratio increased significantly, averaging a 6-fold increase from a single culture (Figure 4B and C).

This selection strategy has several key advantages. By separating the counterselection module into a separate cell, mutations that prevent proper biosensor function yield false negatives rather than false positives, by preventing producers from gaining the expected benefits of muconate production. When selecting for rare phenotypes, false negatives are much more easily tolerated than false positives. Similarly, the competitor strain can be eliminated and reintroduced after every round of growth, preventing the accumulation of strains with mutated biosensors.

This system also streamlines the process of using biosensors in diverse bacteria. The producer and competitor strains can be entirely different species, individually chosen for specific purposes. Rather than engineering the muconate biosensor into *P. putida*, we were able to leave it in the more-tractable *E. coli* host. This muconate-sensitive competitor can then be used with any muconate producing strain that has compatible culture conditions. The producer strain, meanwhile, has no remnant of the biosensor that needs to be removed later. We also note that the biosensor is not strictly necessary, as long as a competitor strain can be identified or engineered that is inhibited by the product of interest. Finally, the validation of a new biosensor is simplified. Frequently, biosensors are validated through exogenous addition of small molecules and then used to sense compounds produced intracellularly. Here, the competitor strain is intended to respond to exogenous products, so testing and application are performed under equivalent conditions.

The selective pressures that result from this approach are complex. An increase in either growth rate or muconate production would be expected to provide a selective advantage. In general, we expect this complex selective pressure to be a benefit of this approach, since the parallel selection for growth rate prevents the accumulation of mutations with strong fitness costs and will yield robust strains. The selective pressure can also be tuned in multiple ways, including modifying the sensitivity of the biosensor to its target, changing the initial population ratio of producers and competitors, or engineering the competitor to grow more or less quickly.

While the initial selective advantage for a single round of growth in droplets was modest, 5.8 ± 1.5-fold, this represents a clear selection against nonproductive mutations. Mutant strains with increased productivity will likely gain an additional selective advantage and produce a larger fold enrichment.

New methods for strain and library construction have expanded the opportunities for large-scale biological design. However, testing the resulting strains currently limits throughput. To address this bottleneck, we have developed a flexible biosensor-driven selection strategy using bacterial competition in microfluidic droplets. Using this approach, libraries generated through a variety of methods can be rapidly tested to identify rare variants that improve productivity. By increasing testing throughput, we can accelerate the development of new production strains and shorten the time to market.

## Materials and Methods

### Media and chemicals

All chemicals were from Sigma-Aldrich unless otherwise noted. Strains were routinely propagated in LB or M9 minimal medium containing 2 g/L glucose. Antibiotics were 50 µg/mL kanamycin, 100 µg/mL ampicillin, and 50 µg/mL streptomycin. Stock solutions of muconate and coumarate at 100 mg/mL were made fresh in DMSO and diluted to the appropriate concentrations as needed. Microfluidic droplets used HFE-7500 (3M, St. Paul, MN) as the bulk phase and 008-FluoroSurfactant (RAN Biotechnologies, Beverly, MA) as the surfactant.

### Genetic manipulation

Strains JMP15 and JMP26 contain multiple mutations relative to the parental strain CJ102. For gene deletions, the appropriate deletion construct, containing approximately 500 bp of homology on both sides of the gene to be deleted, were synthesized *de novo* (IDT, Skokie, IL) and assembled into pK18mobsacB^21^ using the HiFi Assembly Master Mix (NEB, Ipswich, MA). The resulting plasmids were transformed into *P. putida* by electroporation, followed by selection on LB + kanamycin and counterselection on YT + 25% sucrose^22^. Gene deletions were verified by colony PCR and Sanger sequencing (Eurofins Genomics, Louisville, KY). To introduce the streptomycin resistance marker, plasmids pUX-BF13 and pBK-miniTn7-gfp3 were cotransformed into the recipient *P. putida* strain by electroporation, followed by selection on LB + kanamycin.

### Growth curves

Strains for growth curve experiments were grown to saturation overnight at 30 °C in M9 + 2 g/L glucose + ampicillin. The cultures were then diluted 100-fold into fresh medium and grown in 48-well plates in an Epoch 2 plate reader (BioTek, Winooski, VT) at 30 °C. Growth rates were calculated using CurveFitter software^23^.

### Bulk competition assays

Strains JMP15, JMP26, and REL606+pJM242 were grown to saturation overnight at 30 °C in M9 + 2 g/L glucose + ampicillin. The cultures were washed once and then diluted into 5 mL of fresh M9 + 4 g/L glucose + ampicillin + streptomycin containing the relevant substrate as indicated. Inoculation densities were OD 0.01 for JMP15 or JMP26, and 0.3 for REL606+pJM242. For mixed *P. putida* cultures, the inoculation densities for JMP15 and JMP26 were 0.005, and the inoculum cultures were plated on LB + kanamycin as described below to determine the initial population ratio. Cultures were grown overnight at 30 °C, then diluted and plated on LB + kanamycin. For pure culture experiments, colonies were counted and adjusted for the dilution factor. For mixed culture experiments, individual colonies were isolated and genotyped by colony PCR of the *fcs* locus.

### Droplet competition assays

Droplet competition assays were performed in a similar fashion as bulk assays. The *P. putida* inoculation densities were decreased to 0.002 for pure cultures or 0.001 for each mixed culture, to produce an average of one *P. putida* per 10 droplets. The *E. coli* inoculation density was maintained at 0.15, yielding an average of approximately 6 *E. coli* per droplet. Initial population ratios were determined by plating dilutions on LB + kanamycin agar.

Droplet encapsulation was performed on a Dolomite µEncapsulator system using a 50 µm fluorophilic droplet chip, according to the manufacturer’s directions. The *P. putida* and *E. coli* cell suspensions were placed in separate channels of the reservoir immediately before encapsulation. Flow rates of the driving fluids for *P. putida* and *E. coli* were set to 7 µL/min and the surfactant-in-oil solution was held constant at 22 µL/min. Stability of the droplet generation process was visually monitored to validate each run to completion. A typical run generated approximately three million droplets with an average diameter of 60 µm. After one culture reservoir was exhausted, the collection vial was removed, immediately capped, inspected for quality (no visually distinguishable large droplets observed) and emulsion quantity (~500 µL total, 200 µL emulsion layer), and incubated at 30 °C overnight.

After overnight growth, the emulsion was broken through the addition of 200 µL of perfluorooctanol. The mixture was gently mixed by flicking the tube 5 times, then allowed to settle for 5 min. The top aqueous layer (~120 µL) was removed into a new sterile microfuge tube and serial dilutions (10^-1^ to 10^-3^) plated onto LB + kanamycin agar. Following overnight growth at 30 °C, densities were determined through colony counting, and population ratios by colony PCR of the *fcs* locus as described above.

## Tables

**Table 1:**
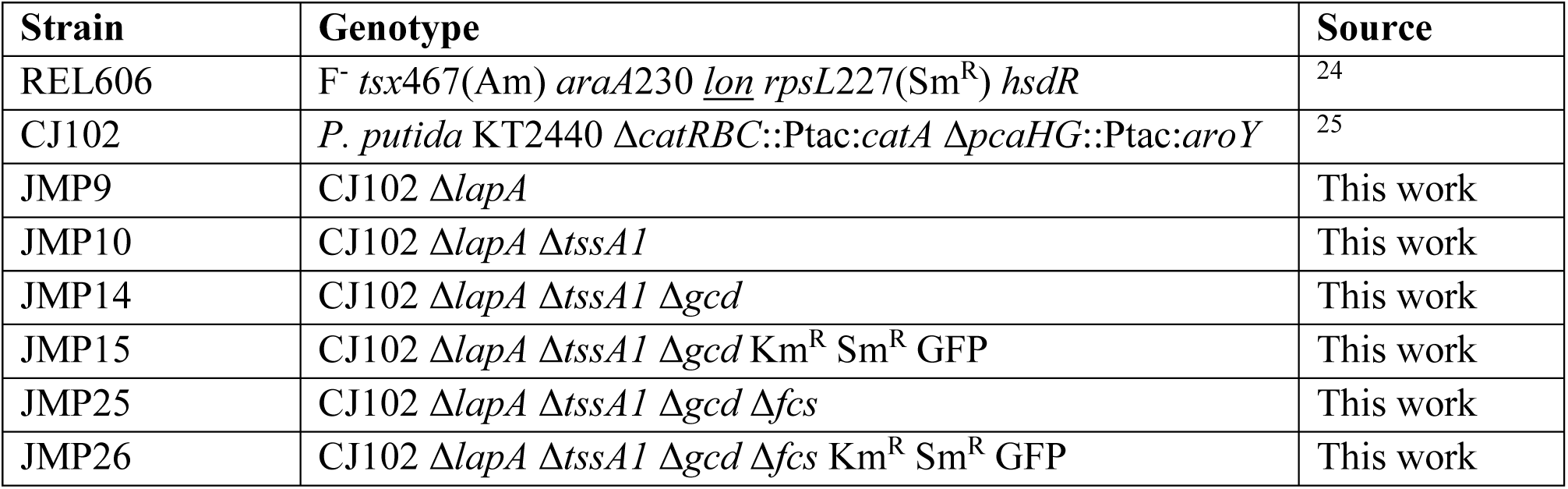
Strains used in this work.

| Strain | Genotype | Source |
| --- | --- | --- |
| REL606 | F<sup>-</sup> *tsx467*(Am) *araA230* *lon* *rpsL227*(Sm<sup>R</sup>) *hsdR* | <sup>24</sup> |
| CJ102 | *P. putida* KT2440 Δ*catRBC::Ptac:catA* Δ*pcaHG::Ptac:aroY* | <sup>25</sup> |
| JMP9 | CJ102 Δ*lapA* | This work |
| JMP10 | CJ102 Δ*lapA* Δ*tssA1* | This work |
| JMP14 | CJ102 Δ*lapA* Δ*tssA1* Δ*gcd* | This work |
| JMP15 | CJ102 Δ*lapA* Δ*tssA1* Δ*gcd* Km<sup>R</sup> Sm<sup>R</sup> GFP | This work |
| JMP25 | CJ102 Δ*lapA* Δ*tssA1* Δ*gcd* Δ*fcs* | This work |
| JMP26 | CJ102 Δ*lapA* Δ*tssA1* Δ*gcd* Δ*fcs* Km<sup>R</sup> Sm<sup>R</sup> GFP | This work |

**Table 2:**
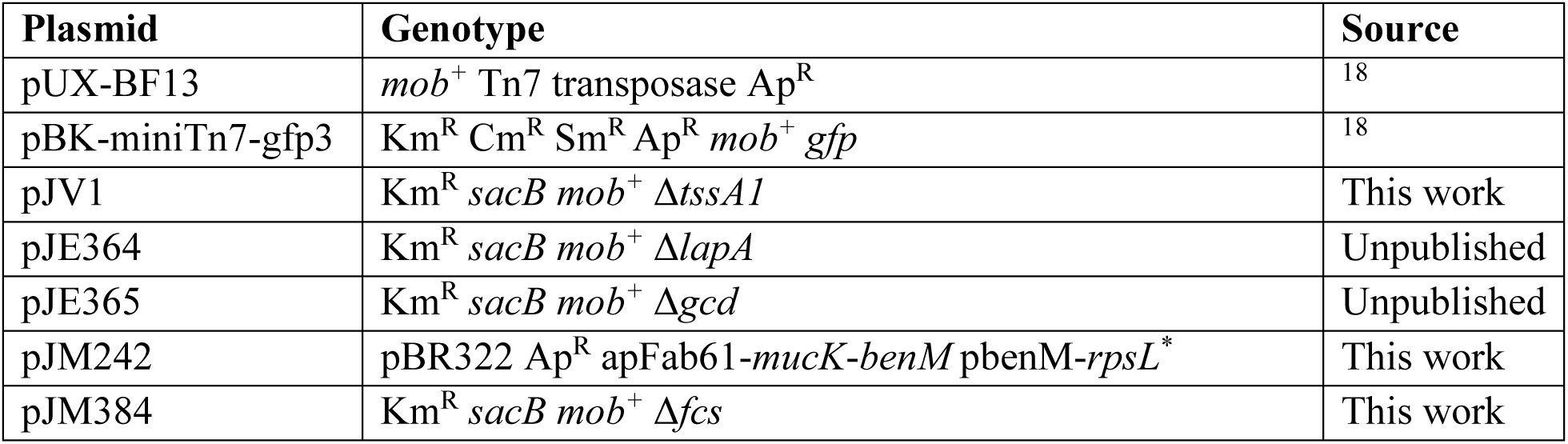
Plasmids used in this work

## Acknowledgements

Gregg Beckham and Chris Johnson graciously provided strain CJ102. Strains pUX-BF13 and pBK-mini-Tn7-gfp3 were provided by Ole Nybroe. Plasmids pJE364 and pJE365 were provided by Joshua Elmore and Adam Guss. Oak Ridge National Laboratory is managed by UT-Battelle, LLC, for the DOE under Contract No. DE-AC05-00OR22725.

